# Increased dietary levels of folic acid reduce survival and alters climbing behaviors 24 hours after hypoxia in male and female *Drosophila melanogaster*

**DOI:** 10.1101/2024.10.23.619884

**Authors:** Siddarth Gunnala, Alisha Harrison, Amber Juba, Paula Ashcraft, Meher Garg, Teodoro Bottiglieri, Lori M Buhlman, Nafisa M. Jadavji

## Abstract

Hypoxia is a major component of ischemic stroke. The prevalence of ischemic stroke is expected to increase as the global population ages and risk factors, like obesity, are on the rise. Nutrition is a modifiable risk factor for ischemic stroke. Increased dietary intake of folic acid (FA) has become an increasing problem in the U.S and other countries as more people are consuming at or above the recommended daily amount of FA. The impact of too much dietary FA on hypoxia is not well understood. This study aimed to investigate how increased dietary levels of FA impact hypoxia outcomes using *Drosophila melanogaster* as a model. Adult female and male *w^1118^ Drosophila melanogaster* flies were placed on control (CD) and 100 µM folic acid (FAD) diets. At 5 to 6 days old fruit fly progeny were exposed to hypoxia for two hours prior to returning to normoxic conditions. We observed escape behavior in hypoxia larvae, confirming exposure to hypoxia. Using liquid chromatography-tandem mass spectrometry, elevated FA levels were observed in FAD compared to controls. We report increased acute hypoxia-induced mortality in FAD flies. Furthermore, FAD flies were not motivated to climb after hypoxia. Under normoxic conditions FAD flies had a higher velocity when descending during a climb. Interestingly, there was no impact of FA on apoptosis in brain tissue post-hypoxia. Together these data suggest that increase dietary intake of FA can have negative health outcomes after hypoxia.

**Highlights:**

- High dietary folic acid and hypoxia increased mortality at the 24-hour time point.
- Hypoxia and folic acid decreased motivation to climb and reduced movement in flies.
- No effect of hypoxia and folic acid on apoptosis levels in the fly brain tissue.

## 1. Introduction

Folic acid (FA) is well known for its role in pregnancy as the demand for it increases significantly. Inadequate levels of FA during pregnancy lead to infant neural tube defects (NTD)[1]. In 1998, the U.S and Canadian governments began fortifying foods with FA to reduce the prevalence of NTDs. Furthermore, the movement towards optimizing health through proper diet and supplementation started to gain popularity. Research followed suit with most published work focusing predominantly on the adverse effects and risks associated with deficiencies in FA [2]. However, the impact of increased levels of FA on other health outcomes is not well characterized.

There have been reports indicating that too much dietary intake of FA can be detrimental to health [3–5]. This includes promotion of tumorigenesis [3,4], negative interactions when levels of B12 are reduced [5] and negative outcomes in offspring when mothers are fed too much folic acid during pregnancy [6]. The Centers for Disease Control recommends 400 µg per day of FA daily for adults [7]. Over-supplementation is defined as a consumption of FA over 1,000 µg per day in adults [8]. A study in the US reported that 54.9% of the sampled population excluding pregnant and lactating women were taking ≥ 400 µg/d of FA [9].

In populations where no mandatory FA fortification is not in place, supplementation with FA has been shown to have positive effects when individuals are at high risk of stroke [10,11]. Results from an *in vitro* study investigating FA supplementation of 4µg/mL prior to hypoxia in human monocyte THP-1 cells showed anti-inflammatory effects, through downregulation of interleukin-1beta (IL-1β) and tumor necrosis factor-alpha (TNF-α) protein levels [12]. In this same study, vascular endothelial growth factor (VEGF) mRNA levels are decreased in a dose-dependent manner in response to FA and hypoxia treatments. Additionally, FA pretreatment attenuates nuclear hypoxia inducible factor-1 alpha (HIF-1α) signal, which could lead to worse chronic hypoxia management due to less angiogenesis [12]. Other *in vitro* studies have shown that FA reduced the inflammatory response by inhibiting reactive oxygen species [13] and increased cell survival [14] post hypoxia. In mice a combination of increased B-vitamins and choline ameliorated hypoxia-induced memory deficit [15]. However, the impact of FA over-supplementation *in vivo* requires more investigation. The objective of our present study is to understand the role of increased dietary intake of FA and hypoxia on survival, larvae behavior, and climbing behaviors *in vivo* using the *Drosophila melanogaster* model.

## 2. Methods

### 2.1. Experimental timeline and *Drosophila melanogaster* model

Our experimental manipulations are summarized in **Figure 1**. Manipulations included placing adult *w^1118^* flies (Bloomington Drosophila Stock Center, Indiana University, Bloomington, IN, USA) on either folic acid diet supplemented (FAD) or control diets for a period ten days. This ten-day period is roughly the amount of time it takes for larval development to finish and allow the offspring to emerge from pupa cases as adult flies [16]. After about 5-6 days post-eclosion, adult flies are either subjected to hypoxia for a period of 2 hours or left in atmospheric oxygen levels (normoxia). Twenty-four hours after the exposure to hypoxia, the flies were subjected one of the following experiments: mortality, climbing assay, larvae escape time, or brains tissue dissection and immunofluorescence. Additionally, non-hypoxia control diet and FAD fly whole bodies were collected for FA and related metabolite measurements. The flies were housed in a controlled laboratory environment, 25ºC with consistent 12-hour light-dark cycles.

**Figure 1.**
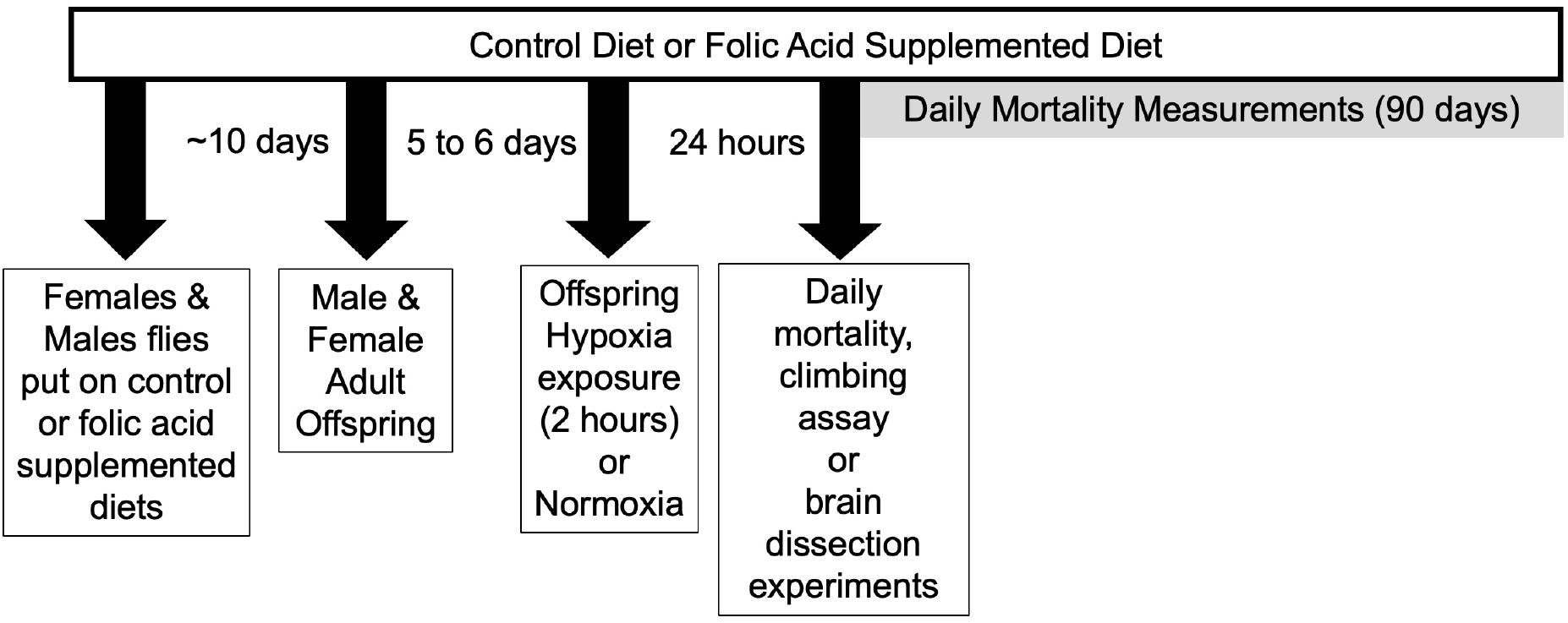
Experimental timeline for experiments. Adult female and male *w^1118^ Drosophila melanogaster* were placed on one of 2 diets: control diet (CD, 0µM folic acid) or folic acid diet (FAD, 100µM folic acid). Offspring from the crosses were used for experimental studies. At 5 to 6 days of age female and male flies were exposed to 2 hours of hypoxia at 1% oxygen or normoxia conditions. After which all flies were maintained in normoxic conditions to model reperfusion. Experimental files were allocated to the following experiments: daily mortality measurements were taken for 90 days post hypoxia. Twenty-four hours post-hypoxia negative geotaxis (climbing) assays, or brain tissue was dissected for immunofluorescence experiments. Other groups of flies were used to measure the larvae escape behavior and folic acid metabolites.

### 2.2 Increasing Dietary Intake of Folic Acid in Fly Food

Standard fly food consisting of a mixture of corn syrup, cornmeal, molasses, agar, soy flour, and yeast was made in the laboratory using standard methods and protocols.

Two dietary conditions were included in the present study, a control diet (CD, 0 µM FA), or a supplemented folic acid diet (FAD) with a final concentration of 100 µM FA (MilliporeSigma, Burlington, MA, USA). Food vials were stored at 4ºC until used. Experimental flies’ vials replaced every 7 days for the duration of experimental studies.

### 2.3 Hypoxia Treatment

An Eppendorf CellXpert (C170i) was used to induce hypoxia in flies. Flies 5-6 days post eclosion were subjected to 1% O_2_, 26°C, and 0.1% CO_2_ for 2 hours. After hypoxia exposure, flies were transferred to normoxic (20% O_2_) conditions for 24 hours to model reperfusion injury [17].

### 2.4 Behavioral Escape Response of Larvae to Hypoxia

To confirm exposure to hypoxia we performed larvae climbing experiments [18,19]. These experiments were performed in the hypoxia chamber, using the transparent cover to count larvae that moved out of food vial as demonstrated in Figure 2A. Vials containing larvae at all stages were placed in the hypoxia chamber via their respective FA treatment vials, along with flashlights to illuminate the interior of the chamber. The chamber was activated and allowed to reach the desired point of hypoxia. At that stage, the number of larvae in each of the 12-29 vials per condition group that exhibit a behavioral escape response was recorded at time 0, 5, and 10 minutes. We measured larvae from control and FAD diets, as well as normoxia and hypoxia conditions.

**Figure 2.**
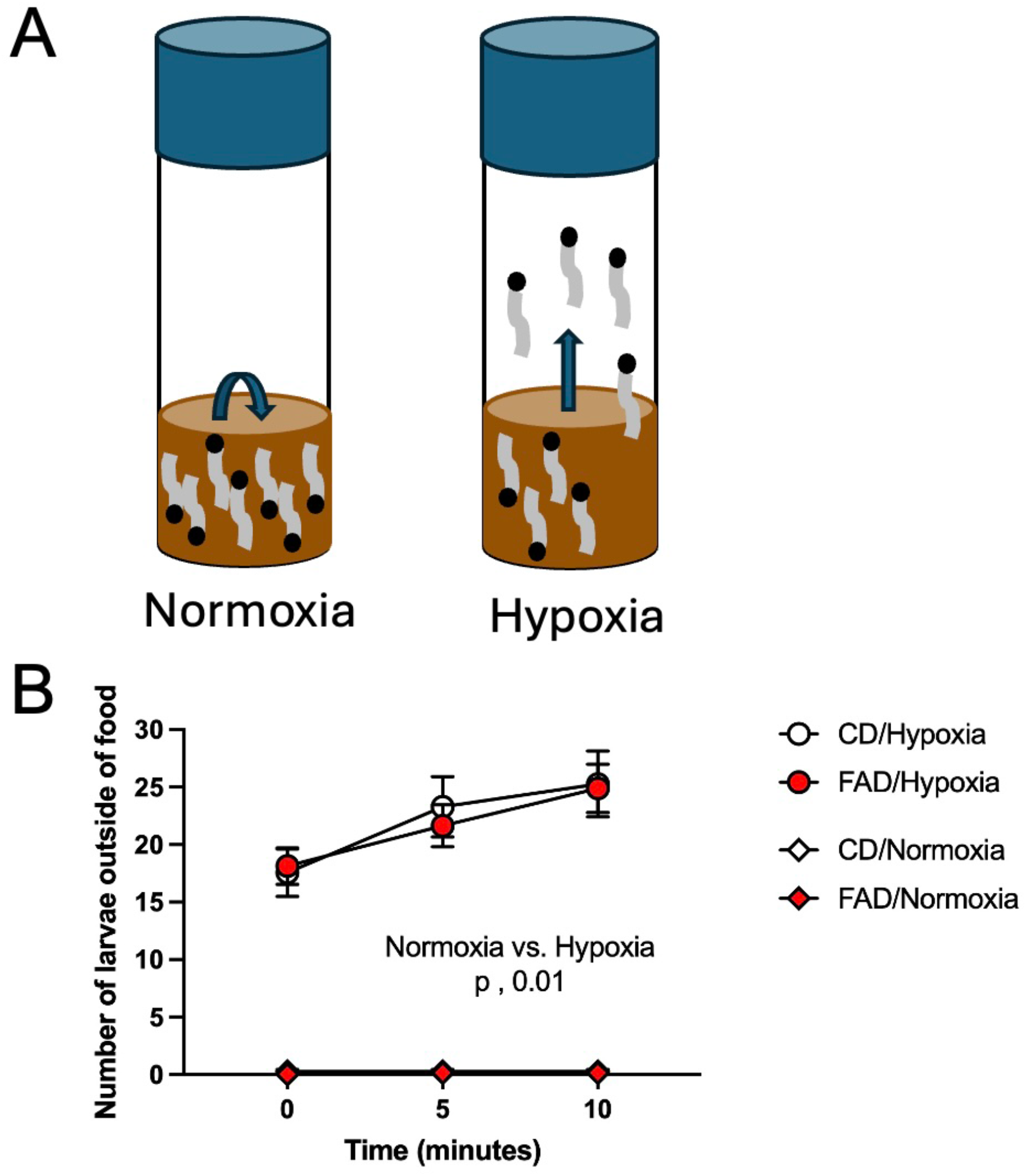
Impact of hypoxia and increased dietary intake of folic acid (FAD) or control (CD) diets on larvae escape behavior. (A) Visual diagram of fly vials depicting differences of exploratory and feeding behaviors for fly larvae in conditions of normoxia and hypoxia. (B) Quantification of the number of larvae escaping at time 0, 5, and 10 after onset of hypoxic conditions (1%) compared to normoxic larvae. There was a significant impact of hypoxia on larvae escape behaviors (p < 0.001). The data represent the mean ± standard error of the mean (SEM) of 12-29 vials per group.

### 2.5 Folic Acid Metabolite Measurements

Six whole body tissue lysates of FA-treated 5–6-day post-eclosion female and male flies were tested. Briefly, an aliquot of 20 mg frozen tissue was transferred to homogenization tubes and were homogenized using a tissue homogenizer. After homogenisation, charcoal-treated rat plasma and centrifuged. After centrifugation, 300 μL of supernatant was passed through a 10-kDa cut-off filter at 14,000 rpm and the filtrate was collected for injection into the chromatographic system for FA, tetrahydrofolic acid (THF), and 5,10-methenyltetrahydrofolate using a UHPLC– MS/MS system consisting of a Waters Acquity UHPLC coupled to a Xevo-TQS triple quadrupole mass spectrometer equipped with an electrospray ionization probe (Waters Corporation, Milford, MA, USA). All calibration curves analyzed during validation were linear for the standards ranging from 0.5 to 2500 ng/mL for FA, tetrahydrofolic acid (THF), and 5,10-methenyltetrahydrofolate [20].

### 2.6 Survival Assay

Starting 24 hours post-hypoxia exposure (n = 204 to 465 individual flies), we recorded the number of individual dead and surviving flies were recorded daily until all flies expired. Approximately 20 to 25 vials of flies from hypoxia and normoxia conditions were used for this assay.

### 2.7 Climbing Assay

We measured climbing behavior in flies to negative geotaxis in flies [21]. The behavioral assay was performed on flies 24 hours after exposure to hypoxia or normoxia. Flies from each experimental group were placed individually into one of 16 transparent 5 mm diameter, 80 mm long polycarbonate tubes. The MB5 Multibeam Activity Monitor (Trikinetics Inc. Waltam, MA, USA) holds 16 polycarbonate tubes in a vertical position with 9 infrared beams passing through each tube at 3 mm vertical distances for a total of 51 mm, leading to a total of 17 unique positions in the tube. The Multi-Beam Monitor (MBM) device records the position of the fly in the tube each second for 20-minutes, counting a movement as any positional change that was captured per second through singular or multiple beam breaks. Every time the fly ascends to a peak and to reverse travel for a minimum of 3 mm. The total upward movement is determined by counting the number of beam breaks as a fly ascends. Individual fly behavior was measured; there were 14 to 29 flies used per experimental group.

### 2.8 Immunofluorescence

Brain tissue from 9 to 14 flies was used for immunofluorescence analysis to assess molecular mechanisms involved hypoxia and increased dietary intake of FA. Brain tissue was fixed with using 3.7% formaldehyde and then blocked using normal goat serum (NGS, 1:10) diluted in PBS-T. The active caspase-3 primary antibody (ac3; 1:100, Cell Signaling Technologies, Danvers, MA, USA, Catalog: # 9662S, RRID: AB_331439) was diluted in PBT and incubated with brain tissue overnight at 4°C. The next day, whole brains were incubated in an anti-rabbit Alexa Fluor 555-conjugated secondary antibody (1:100, Cell Signaling Technologies Danvers, MA, USA, Catalog: 4413S) and then mounted for microscope analysis. Z-stacks of active caspase-3 per brain were captured at 400X magnification with a TCS PSE confocal microscope (Lecia Microsystems, Wetzlar, Germany) and analyzed with Image Pro Premier 3D image processing software (Media Cybernetics, Inc., Rockville, USA). All microscope procedures were conducted by individuals blinded to experimental groups.

### 2.9 Statistics

The survival curve after exposure to hypoxia or normoxia with varying levels of FA was analyzed using a Hillslope curve. This test utilized probability curves derived from an application of the Hill equation on the control flies using the HEPB (Hill Equation with Probability Band) program [22]. This analysis provided the predicted day at which each condition’s group’s population will have a 50% life expectancy (LE50). The LE50 value of each condition group was then compared to the true control being flies kept in normoxic conditions with a control diet. A 95% prediction interval for the confidence band was applied to the data based on an algorithm by N.C Shammas that approximates critical values in the distribution.

Using GraphPad Prism 10 we performed the D’Agostino-Pearson normality test prior to t-test and ANOVA testing. The larvae escape time was analyzed using two-way ANOVA with repeated measures. Climbing attempts, total height climbed, descending velocity, and percent expired flies on day 1 post-hypoxia were analyzed using a three or two-way ANOVA and significant main effects were followed up with Tukey’s multiple comparisons test. Active-caspase 3 immunofluorescence volume data was quantified using an un-paired t-test via GraphPad Prism 10. The mean and the standard error of the mean of each group is presented on the graphs unless otherwise indicated.

## 3.0 Results

### 3.1 Larval Escape from Food During Hypoxia

The escape behavior of larvae from food occurs under hypoxic conditions [18,19]. We used the larvae escape behavior to confirm that our flies were subjected to hypoxia in the incubator we were using. A cartoon and representative image of larvae under normoxia and hypoxia conditions is shown in **Figure 2A**. When exposed to hypoxia, larvae from both CD and FAD groups crawled up out of the food when compared to normoxic larvae (**Figure 2B**; F(_1,57_) = 187.88, *p* < 0.001). There was no effect of diet (F(_1,57_) = 0.25, *p* = 0.62), and there was no interaction between hypoxia and diet (F(_1,57_) = 0.16, *p* = 0.69). We did not observe larvae escape from food in vials under normoxic conditions.

### 3.2 Increased Levels of Whole-Body Folic Acid in FAD Flies

To confirm over increased levels of FA, we measured folic acid related metabolites in CD and FAD whole bodies of flies. For these set of experiments, we measured levels of metabolites in flies that were not exposed to hypoxia (normoxic flies). We measured FA, tetrahydrofolic acid (THF), and 5,10-methylenetetrahydrofolate (5,10-CH_2_-THF) levels using mass spectrometry. FA levels were elevated in female and male FAD flies compared to CD (**Figure 3A**; F(_1,20_) = 122.1, *p* < 0.0001). There was no effect of sex (F(_1,20_) = 1.70, *p* = 0.21) or interaction between FA levels and sex (F(_1,20_) = 1.83, *p* = 0.1913).

**Figure 3.**
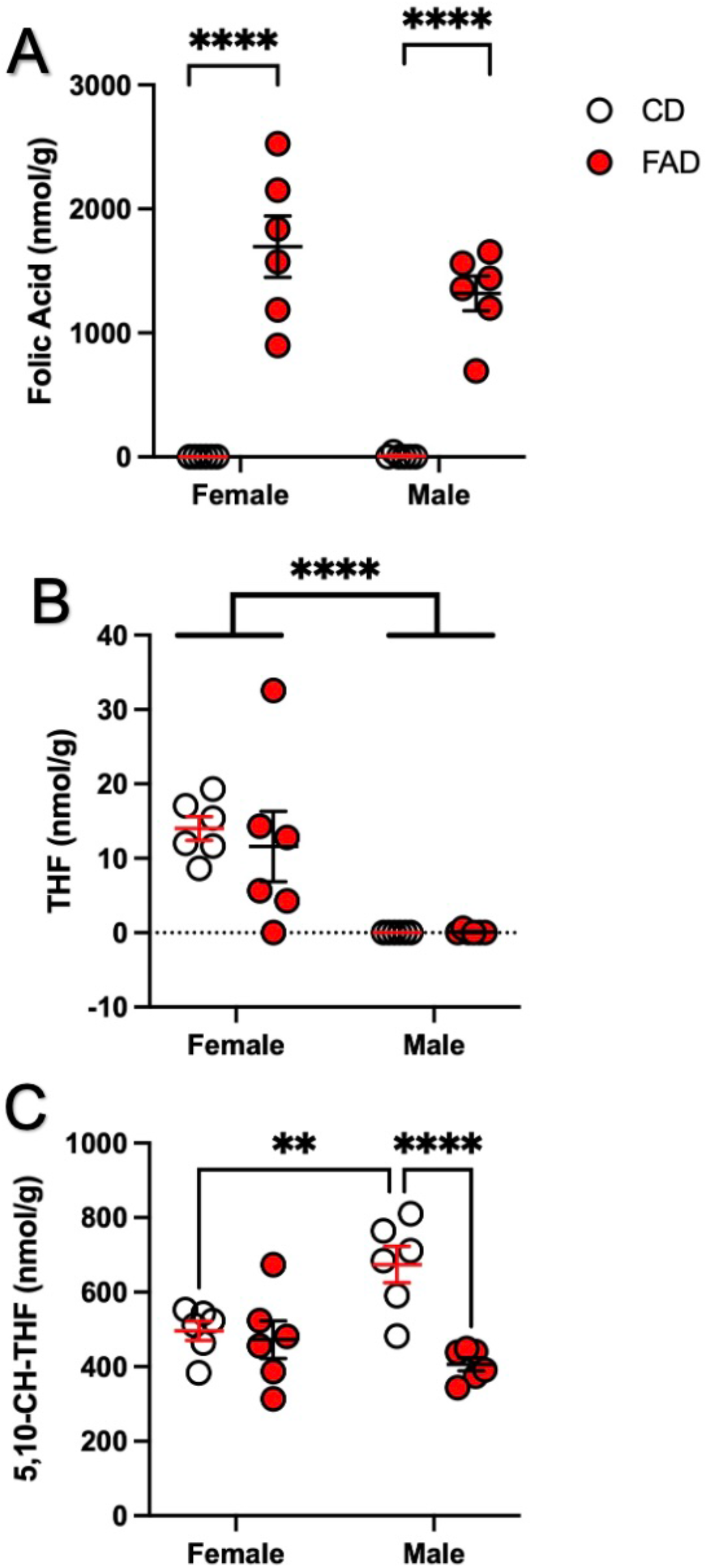
Impact of increased dietary folic acid (FAD) or control diet (CD) on whole body folic acid (A), Tetrahydrofolate (THF) (B), and 5,10-CH-THF (C) content in flies maintained on control or 100µM folic acid supplemented diets. The data represent mean ± standard error of mean (SEM) of 6 flies per group. A,C ** p < 0.01, **** p < 0.0001 Tukey’s pairwise comparisons. B, **** p < 0.0001 2-way ANOVA sex main effect.

Female flies of both control diet and FAD had higher THF levels (**Figure 3B**; F(_1,20_) = 26.03, *p* < 0.0001; control, *p* = 0.0039, FAD, *p* = 0.02). Diet had no effect on THF levels (F(_1,20_) = 0.2140, *p* = 0.65), and no interaction between FA levels and sex were detected (F(_1,20_) = 0.25, *p* = 0.62).

CD flies had higher levels of 5,10-CH_2_-THF (**Figure 3C**; F(_1,20_) = 14.33, *p* = 0.001). This effect was driven by males (p < 0.0001). Untreated female flies had reduced levels of 5,10-CH_2_-THF compared to untreated males (*p* = 0.005). There was a significant interaction between FA levels and sex (F(_1,20_) = 10.07, *p* = 0.005), and there was no effect of sex on THF levels (F(_1,20_) = 2.098, *p* = 0.16).

### 3.3 Reduced FAD Fly Survival 24 hours After Hypoxia

Beginning twenty-four hours after hypoxia treatment, the percent of surviving flies was recorded for 90 days. The 50% life expectancy (LE_50_; is the day at which 50% mortality occurred). This value for the control/normoxia flies was 39 days, while the LE_50_ for FAD/normoxia flies was 45 days **(Figure 4A).** The Hillslope curve LE_50_ was determined utilizing the 95% prediction interval upper and lower probability bands from the control/normoxia data. LE_50_ for upper and lower bands are 47 and 30 days, respectively **(Figure 4A)**. LE_50_ values less than 30 days are considered statistically significant [22]. The LE_50_ control/hypoxia was 8 days (*p* < 0.05), FAD/normoxia was 45 days, FAD/hypoxia 5 days (*p* < 0.05) **(Figure 4A)**.

**Figure 4.**
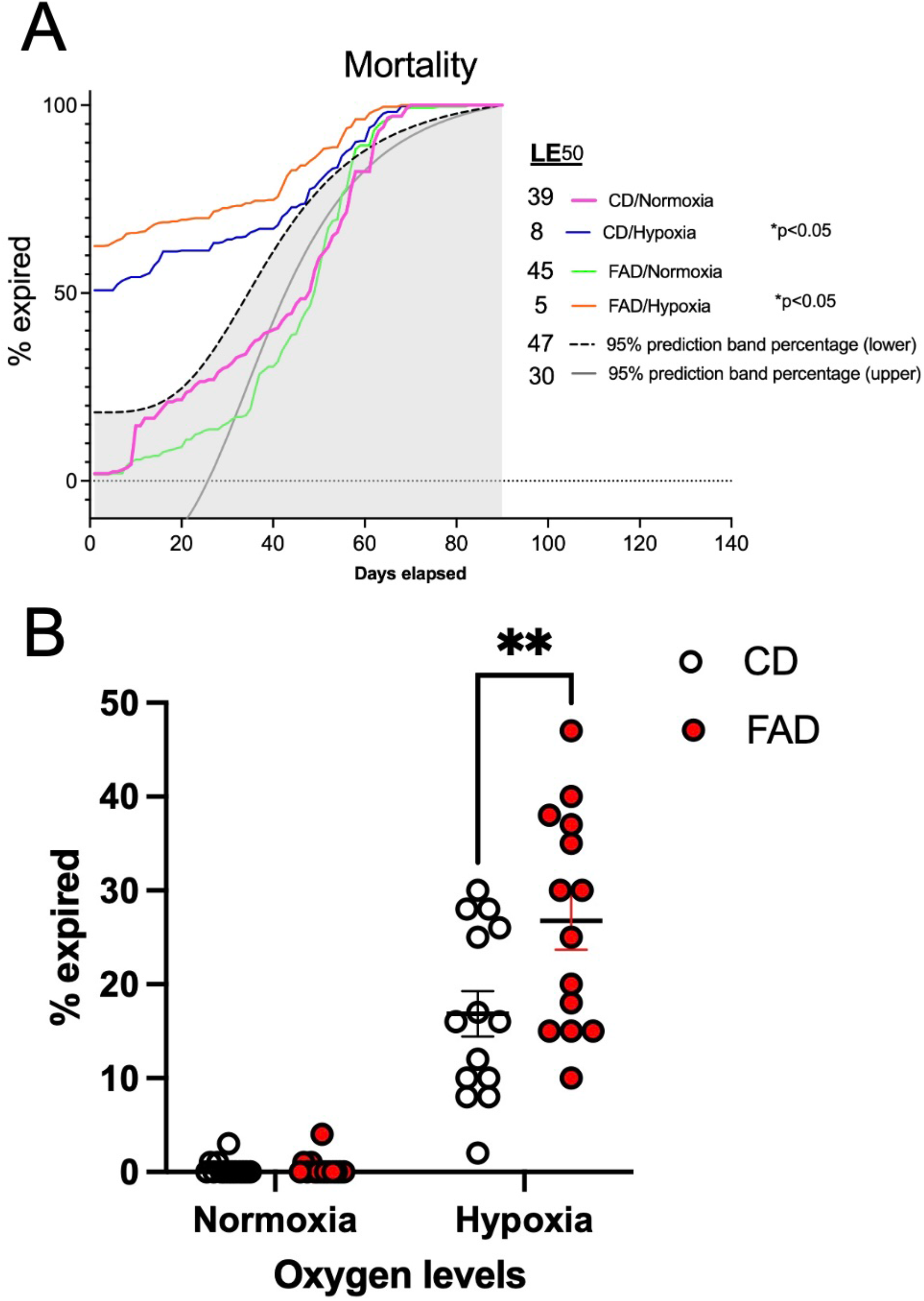
Survival study of flies maintained on control (CD) and 100µM folic acid (FAD) subjected to 2 hours of hypoxia or normoxia. (A) Ninety-day percent expired flies. (B) Percentage of expired flies 24 hours after hypoxia. * p < 0.05, ****p < 0.00001, Tukey’s pairwise comparison between indicated groups. The data represent the survival of flies over two studies of 204 to 456 flies. For B, the data represent the mean ± standard error of the mean (SEM) of 12-29 vials per group.

Acute mortality in response to hypoxia was observed the first 24 hours after hypoxia. FAD flies had increased mortality (**Figure 4B;** F(_1,52_) = 6.46, *p* = 0.01) Additionally there was a significant interaction between fly groups in different FA and oxygen levels (F(_1,52_) = 6.27, *p* = 0.015), and this effect was exacerbated in as well as flies exposed to hypoxia groups in different oxygen levels (F(1,52) = 118.6, *p* < 0.0001). Tukey’s multiple comparisons test demonstrated a difference between control diet normoxia and hypoxia flies (*p* < 0.0001) as well as a difference between FAD normoxia and hypoxia flies (*p* < 0.0001). There was a difference between CD hypoxia flies and FASD hypoxia flies (*p* = 0.004).

### 3.4 Reduced Climbing Behaviors in Flies Post-Hypoxia

Twenty-four hours after hypoxia treatment, climbing behavior was measured in hypoxic and normoxic male and female flies. There were two specific measurements we made, the first was the total movement and the second was the motivation to climb.

#### 3.4.1 Total Movement

Data with both male and female flies combined because no sex difference was observed (F(_1,139_) = 0.39, *p* = 0.53). Flies exhibited a significant difference between hypoxia and normoxia in the number of total movements (**Figure 5A**; F(_1,143_) = 4.76, *p* = 0.031). There was no effect of FA (F(_1,143_) = 0.65, *p* = 0.42) or any interaction between fly groups of different FA and oxygen levels (F(_1,143_) = 0.77, *p* = 0.38).

**Figure 5.**
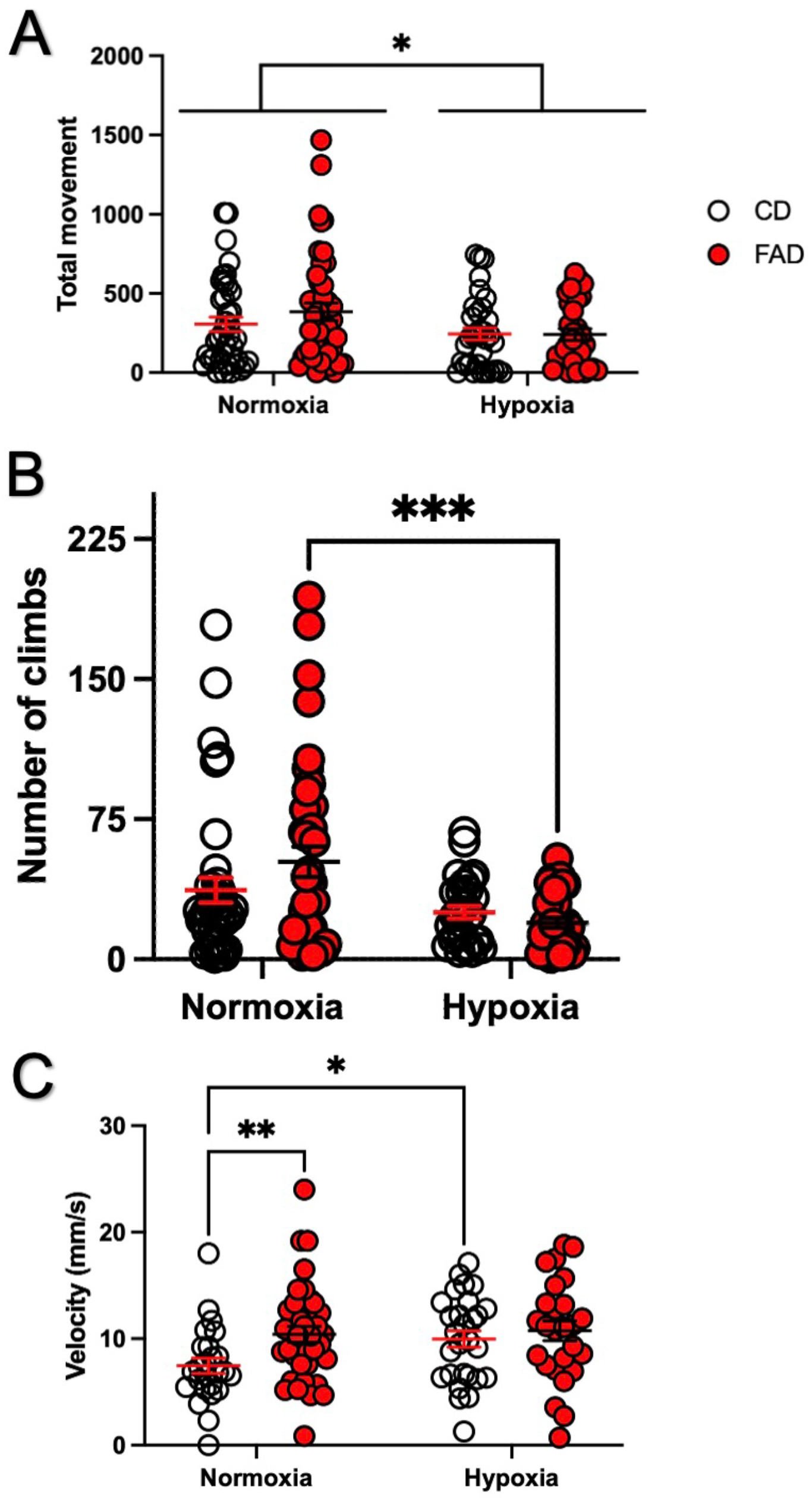
Impact of hypoxia and increased dietary intake of folic acid (FAD) or control diet (CD) on fly climbing behaviors. (A) Total climbing movement, * p = 0.031 oxygen treatment main effect. (B) Motivation to climb (number of climbs), *** p = 0.005 Tukey’s pairwise comparison. (C) Descending velocity, * p = 0.036, ** p = 0.0087 Tukey’s pairwise comparison. The data represents the mean ± standard error of the mean (SEM) of 29 to 41 flies per group.

#### 3.4.2 Motivation to Climb

There were no differences between female and male flies (F(_1,129_) = 1.22, *p* = 0.27), so they have been grouped together. CD and FAD flies exposed to hypoxia had reduced motivation to climb, as indicated by number of climbs (**Figure 5B**; F(_1,133_) = 12.60, *p* = 0.001). There was no effect of FA (F(_1,133_) = 0.58, *p* = 0.45) or interaction between hypoxia and diet (F(_1,133_) = 2.76, *p* = 0.09). Tukey’s multiple pairwise comparisons demonstrated an interaction between normoxia and hypoxia FAD flies (*p* = 0.001), as well as an interaction between FAD normoxia and CD hypoxia flies (*p* = 0.02)

#### 3.4.3 Descending Velocity

There were no differences between female and male flies (F(_1,101_) = 2.53, *p* = 0.061) in their descending velocity measurements, so they have been grouped together. Flies exhibited a significant difference between FAD and CD flies in their descending velocity (**Figure 5C**; F(_1,113_) = 5.32, *p* = 0.023). Normoxia FAD flies had a higher velocity than CD flies (p = 0.087). Hypoxic CD flies had a higher velocity compared to normoxia CD flies (p = 0.0036). There was no effect of oxygen levels (F(_1,113_) = 0.65, *p* = 3.08 or any interaction between fly groups of different FA and oxygen levels (F(_1,113_) = 1.86, *p* = 0.18).

### 3.5 No Changes in Apoptosis in Brain Tissue of FAD flies After Hypoxia

We measured apoptosis in female and male flies that were exposed to hypoxia in brain tissue. These flies were maintained on FAD and control diets. Due to differences in brain size between male and female brains, active-caspase 3 volume was quantified separately between hypoxia between the sexes. Unpaired t-test of male (**Figure 6A**, *p* = 0.70) and female (**Figure 6B**, *p* = 0.84) brains separately demonstrated no significant differences between control diet and FAD.

**Figure 6.**
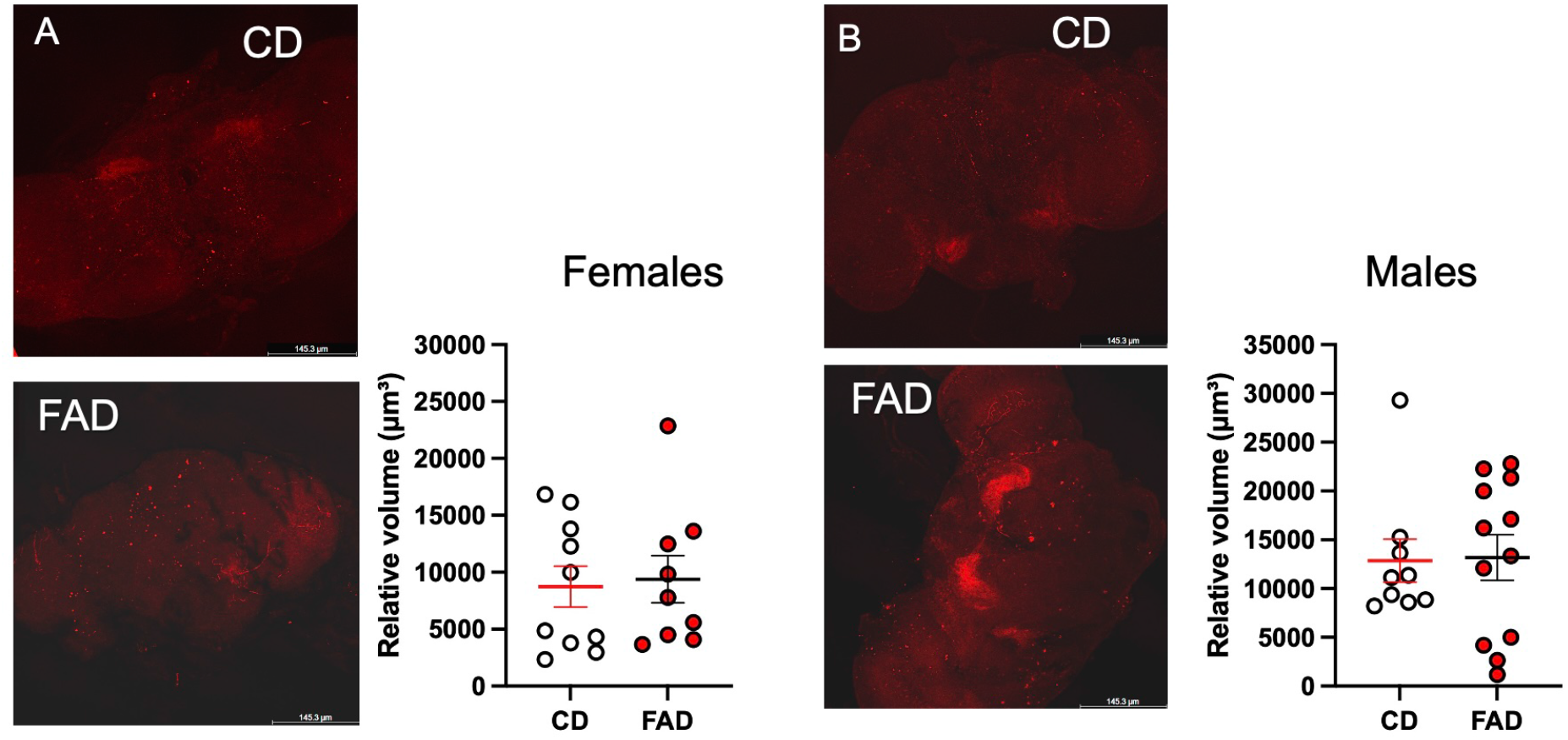
Relative volume of active caspase-3 in brain tissue of control diet (CD) and increased dietary intake of folic acid (FAD) flies 24 hours after exposure to hypoxia, A, females and B, males. The data represents the mean ± standard error of the mean (SEM) of 9 to13 brains used per diet group.

## 4.0 Discussion

Hypoxia is a major component of ischemic stroke. Nutrition is a modifiable risk factor for ischemic stroke. In countries with mandatory fortification there has been an increase in over supplementing their diets with FA. The impact of too much dietary FA intake on hypoxia, a component of ischemic stroke, is not well understood. A dose of 100 µM of folic acid was chosen for this study. Prior experiments using 50 µM or 100 µM of folic acid combined with hypoxia demonstrated high fly mortality using the 100 µM dose (data not shown).

This study highlights interactions between hypoxia and increased dietary intake of FA *in vivo*. We have demonstrated that increased levels of dietary FA exacerbate impact of hypoxia on mortality at the 24-hour time point. Hypoxia and FA also had a negative effect on climbing behavior, flies had a decreased motivation to climb, and reduced movement. Normoxic conditions in FA increased descending velocity, as well as hypoxic CD. We did not observe an effect of hypoxia and FAD on active-caspase 3 levels in the fly brain tissue. In future studies we plan to incorporate other markers of neurodegeneration. For example, hypoxia inducible factor 1 alpha (HIF-1α) is a transcription factor that is turned on after exposure to hypoxia. It has been thought to organize the cells’ response to hypoxia by regulating angiogenesis, cell survival and other vital processed [23]. Another marker is Nrf-2 (Nuclear factor erythroid 2-related factor 2), which is transcription factor involved in the oxidative stress signaling cascade [24].

In our study, we confirmed that the flies were consuming very high amounts of FA from their diet by measuring FA metabolites in whole body tissue. However, we were unable to determine FA body distribution, or how much was absorbed through the gut. Adding 1C enzymatic data or isolated measurements of these metabolites from fly heads, for example, would help narrow down possible explanations. There are notable sex differences in levels of 1C metabolites like tetrahydrofolate and 5,10-CHF-THF. The lack of differences in THF levels may imply enzyme function leading to THF at this level of 1C metabolism is less affected by FA levels and maybe more influenced by sex. Increased levels of FA impact developing organism as demonstrated in mouse studies [6,25,26] as well as older individuals [27]. Cancer has also been reported to increase as a result of increased dietary FA intake [4,28]. FA dietary deficiencies cause hippocampal changes in a mouse model [29], demonstrating sex differences in an invertebrate model, *Drosophila melanogaster,* in conditions of increased dietary FA rather than deficiency.

In our present study, we observed that the highest mortality per day occurred within 24 hours of hypoxia. We found that FAD flies had increased mortality compared to control diet hypoxia flies. There was no impact of FAD on mortality in normoxic or hypoxic groups throughout the remainder of the survival study. The sheer stress of hypoxic cell damage occurred contributed to increased acute mortality in FAD flies. Increase FA levels in hypoxic conditions has been reported to attenuate IL-10, anti-inflammatory cytokine, levels *in* vitro [12]. An inadequate response to hypoxic cell damage due to attenuation of hypoxia response mechanisms could be demonstrated *in vivo* via increased mortality. The relatively indifferent mortality rates after the first twenty-four hours could be due to robust compensation from the cellular damage that occurs after hypoxia exposure. Prior research has demonstrated such novel brain regions in fly brain, namely regions of the brain that interact with the ventral nerve cord and femoral chordotonal organs that function as proprioceptors for locomotion in *Drosophila melanogaster* [30,31]. Mechanistic insights of how FA diet impacts gross development could be involved in the study of the ventral nerve cord after hypoxia. Furthermore, *Drosophila melanogaster* are relatively resilient to hypoxia [32], and fine-tuning the duration and strength of hypoxia for a could cause sub-lethal effects of hypoxia, and trigger adaptive compensation in behavioral and metabolic metrics [32–35].

To dissect the impact of FA and hypoxia on mortality, an additional study that separates male and female *Drosophila melanogaster* to measure the impact of sex on mortality would provide additional insight, as we and others have described sex differences [29,36–38]. A sex difference study in flies would not only provide data on differences of adaptability to FA over-supplementation after hypoxia between sexes, but it would also show what relative age points in male and female flies are most affected by chronic increased levels of FAD. The results of that potential study would be valuable to corroborate with the 1C metabolite data towards finding potential implications of 1C metabolism aberrancies leading to the exacerbations of mortality. This could potentially imply that there are different risks of hypoxic cell damage exacerbation caused by increased dietary of FA levels between males and females, potentially at different ages in their life cycles.

It is unclear whether strong metabolic support for 1C metabolism occurs in neurons associated with proprioception and locomotion. Indeed, suppression of 1C metabolism enzymes, serine hydroxymethyltransferase-2 (SHMT2) and NAD-dependent methylenetetrahydrofolate dehydrogenase-methenyltetrahydrofolate cyclohydrolase (NMDMC) resulted in motor impairment and mitochondrial defects in flies [39]. Measurement of 1C enzymes in our model would provide insight into how they are utilized in cellular maintenance for neuronal cells associated with locomotion and proprioception. This data could be used to compare the response of increased FA levels to other cells of various denominations of importance for the fly’s ability to survive.

Several studies have highlighted the values of flies as a model for disease of aging [40]. While ischemic stroke can occur at different age groups, there is significantly higher prevalence of it occurring in older demographics [41]. The present study used day 5 to 6 old flies which do not model the current clinical demographics of ischemic stroke. It is important to acknowledge when trying to study such a disease through models. The mortality of flies may be different if exposure to hypoxia occurs later in the fly life cycle. If the present study were to have used older flies, the hypoxia response mechanisms may not be as robust and could have lasting effects throughout the remainder of the fly life cycle, leading to potential effects demonstrated in climbing behavior. It would be valuable to perform longitudinal neurodegeneration measurements at different time points through the adult *Drosophila melanogaster* life span into old age. That data could be relatable to understanding how *Drosophila melanogaster* aging manages aberrances of a high FA diet and exposure to hypoxia.

As the basic science field and public health begin to progressively see more demographics come under the conditions of nutritional over-supplementation, our research will hopefully provide avenues for future research to better understand what the impact of chronic dietary increase of FAD has on neurological diseases *in vivo*.

## Acknowledgements

None

## Abbreviations

1C: one-carbon
CD: control diet
FA: folic acid
FAD: folic acid supplemented diet
IL-1β: interleukin-1beta
TNF-α: tumor necrosis factor-alpha protein levels
VEGF: vascular endothelial growth factor
HIF-1α: hypoxia inducible factor-1 alpha

## Ethics Declarations

Not applicable

## CrediT authorship contribution statement

Siddarth Gunnala: data curation, investigation, methodology, writing-original draft, writing-review and editing. Alisha Harrison: investigation, methodology, writing-review and editing. Paula Ashcraft: methodology. Meher Garg: methodology. Teodoro Bottiglieri: investigation, methodology, writing-review and editing. Lori M Buhlman: investigation, supervision, writing-review and editing. Nafisa M Jadavji: conceptualization, data curation, formal analysis, methodology, project administration, resources, supervision, validation, visualization, writing-original draft, writing-review and editing.

## Funding Sources

None

## Data Availability Statement

Data from study is available on Dryad. http://datadryad.org/share/h06Lh_tyUvRVH2oMV6nzIUohjVulXBNuV1BOaW0d5AM

## Notes

### Competing Interest Statement

The authors have declared no competing interest.

### Summary of Updates

We have revised the manuscript for clarity.

## References

[1] J.D. Van Gool, H. Hirche, H. Lax, L. De Schaepdrijver, Folic acid and primary prevention of neural tube defects: A review, Reproductive Toxicology 80 (2018) 73–84. 10.1016/j.reprotox.2018.05.004.

[2] A.L. Boyles, E.A. Yetley, K.A. Thayer, P.M. Coates, Safe use of high intakes of folic acid: research challenges and paths forward, 74 (2016) 469–474. 10.1093/nutrit/nuw015.

[3] K.C. Strickland, N.I. Krupenko, S.A. Krupenko, Molecular mechanisms underlying the potentially adverse effects of folate, Clin Chem Lab Med 51 (2013) 607–616. 10.1515/cclm-2012-0561.

[4] C.M. Ulrich, J.D. Potter, N. Ulrich, Folate supplementation: Too much of a good thing?, Cancer Epidemiology Biomarkers and Prevention 15 (2006) 189–193. 10.1158/1055-9965.EPI-152CO.

[5] G.E. Mullin, Folate: is too much of a good thing harmful?, Nutrition in Clinical Practice: Official Publication of the American Society for Parenteral and Enteral Nutrition 26 (2011) 84–7. 10.1177/0884533610392238.

[6] R.H. Bahous, N.M. Jadavji, L. Deng, M. Cosín-Tomás, J. Lu, O. Malysheva, K.-Y. Leung, M.-K. Ho, M. Pallás, P. Kaliman, N.D.E. Greene, B.J. Bedell, M.A. Caudill, R. Rozen, High dietary folate in pregnant mice leads to pseudo-MTHFR deficiency and altered methyl metabolism, with embryonic growth delay and short-term memory impairment in offspring, Human Molecular Genetics 26 (2017) 888–900. 10.1093/hmg/ddx004.

[7] CDC, Folic Acid: Sources and Recommended Intake, Folic Acid (2025). https://www.cdc.gov/folic-acid/about/intake-and-sources.html (accessed June 30, 2025).

[8] B.J. Merrell, J.P. McMurry, Folic Acid, in: StatPearls, StatPearls Publishing, Treasure Island (FL), 2024. http://www.ncbi.nlm.nih.gov/books/NBK554487/ (accessed November 15, 2024).

[9] C.M. Pfeiffer, M.R. Sternberg, Z. Fazili, E.A. Yetley, D.A. Lacher, R.L. Bailey, C.L. Johnson, Unmetabolized folic acid is detected in nearly all serum samples from US children, adolescents, and adults., The Journal of Nutrition 145 (2015) 520–31. 10.3945/jn.114.201210.

[10] G.J. Hankey, B vitamins for stroke prevention, Stroke and Vascular Neurology 3 (2018) 51–58. 10.1136/svn-2018-000156.

[11] Y. Huo, J. Li, X. Qin, Y. Huang, X. Wang, R.F. Gottesman, G. Tang, B. Wang, D. Chen, M. He, J. Fu, Y. Cai, X. Shi, Y. Zhang, Y. Cui, N. Sun, X. Li, X. Cheng, J. Wang, X.C. Yang, T. Yang, C. Xiao, G. Zhao, Q. Dong, D. Zhu, X. Wang, J. Ge, L. Zhao, D. Hu, L. Liu, F.F. Hou, K. Cao, L. Chen, P. Gao, R. Gao, X. Ji, N. Li, C. Ma, W. Wang, X. Yang, X. Dai, F. Fan, X. Gao, R. Hui, H. Jiang, J. Jiang, X. Jiang, W. Kong, B. Liu, G. Sun, L. Sun, B. Wang, D. Yin, W. Yang, H. Zhang, C. Zhang, L. Zhao, L. Wang, Y. Chen, A. Huang, Y. Li, J. Wang, R. Xie, C. Yao, D. Zhao, Z. Zhao, L.J. Wei, F. Chen, J. Dong, J. Gu, J. Guo, L. Shen, W. Zhang, Z. Zhang, Z. Wang, X. Li, Y.L. Wan, G. Hu, J. Tan, S. Dong, C. Tong, H. Xu, J. Liu, J. Xu, D. Li, C. Tong, H. Li, C. Liu, J. Xu, T. Zhou, J. Zhou, C. Wu, C. Liu, Y. Wang, L. Xu, Z. Zhou, Z. Zhou, S. Liu, C. Tang, Z. Zhang, J. Tang, Z. Xu, Z. Xiong, Q. Zhu, P. Wu, Efficacy of folic acid therapy in primary prevention of stroke among adults with hypertension in China: The CSPPT randomized clinical trial, JAMA - Journal of the American Medical Association 313 (2015) 1325–1335. 10.1001/jama.2015.2274.

[12] X. Huang, Z. He, X. Jiang, M. Hou, Z. Tang, X. Zhen, Y. Liang, J. Ma, Folic Acid Represses Hypoxia-Induced Inflammation in THP-1 Cells through Inhibition of the PI3K/Akt/HIF-1α Pathway, Plos One 11 (2016) e0151553–e0151553. 10.1371/journal.pone.0151553.

[13] J. Ma, X. Zhen, X. Huang, X. Jiang, Folic acid supplementation repressed hypoxia-induced inflammatory response via ROS and JAK2/STAT3 pathway in human promyelomonocytic cells, Nutr Res 53 (2018) 40–50. 10.1016/j.nutres.2018.03.007.

[14] F. Cheng, J. Lan, W. Xia, C. Tu, B. Chen, S. Li, W. Pan, Folic Acid Attenuates Vascular Endothelial Cell Injury Caused by Hypoxia via the Inhibition of ERK1/2/NOX4/ROS Pathway, Cell Biochem Biophys 74 (2016) 205–211. 10.1007/s12013-016-0723-z.

[15] L. Yu, Y. Chen, W. Wang, Z. Xiao, Y. Hong, Multi-Vitamin B supplementation reverses hypoxia-induced tau hyperphosphorylation and improves memory function in adult mice, Journal of Alzheimer’s Disease 54 (2016) 297–306. 10.3233/JAD-160329.

[16] M.A. Fernández-Moreno, C.L. Farr, L.S. Kaguni, R. Garesse, Drosophila melanogaster as a Model System to Study Mitochondrial Biology, Methods Mol Biol 372 (2007) 33–49. 10.1007/978-1-59745-365-3_3.

[17] K. Mandalaneni, A. Rayi, D.V. Jillella, Stroke Reperfusion Injury, in: StatPearls, StatPearls Publishing, Treasure Island (FL), 2025. http://www.ncbi.nlm.nih.gov/books/NBK564350/ (accessed September 2, 2025).

[18] J.A. Wingrove, P.H. O’Farrell, Nitric oxide contributes to behavioral, cellular, and developmental responses to low oxygen in Drosophila, Cell 98 (1999) 105–114. 10.1016/S0092-8674(00)80610-8.

[19] D.B. Morton, Behavioral responses to hypoxia and hyperoxia in Drosophila larvae: Molecular and neuronal sensors, Fly 5 (2011) 119–125. 10.4161/fly.5.2.14284.

[20] J. Nandania, M. Kokkonen, L. Euro, V. Velagapudi, Simultaneous measurement of folate cycle intermediates in different biological matrices using liquid chromatography–tandem mass spectrometry, Journal of Chromatography B 1092 (2018) 168–178. 10.1016/j.jchromb.2018.06.008.

[21] S.T. Madabattula, J.C. Strautman, A.M. Bysice, J.A. O’Sullivan, A. Androschuk, C. Rosenfelt, K. Doucet, G. Rouleau, F. Bolduc, Quantitative Analysis of Climbing Defects in a Drosophila Model of Neurodegenerative Disorders, J Vis Exp (2015) 52741. 10.3791/52741.

[22] S.R. Gadagkar, G.B. Call, Computational tools for fitting the Hill equation to dose-response curves, J Pharmacol Toxicol Methods 71 (2015) 68–76. 10.1016/j.vascn.2014.08.006.

[23] K.J. Banasiak, G.G. Haddad, Hypoxia-induced apoptosis: Effect of hypoxic severity and role of p53 in neuronal cell death, Brain Research 797 (1998) 295–304. 10.1016/S0006-8993(98)00286-8.

[24] N.M. Jadavji, J. Emmerson, W.G. Willmore, A.J. MacFarlane, P. Smith, B-vitamin and choline supplementation increases neuroplasticity and recovery after stroke, Neurobiology of Disease 103 (2017) 89–100.

[25] D. Wiens, M.C. Desoto, Is high folic acid intake a risk factor for autism?—a review, Brain Sciences 7 (2017). 10.3390/brainsci7110149.

[26] L.K. Murray, M.J. Smith, N.M. Jadavji, Maternal oversupplementation with folic acid and its impact on neurodevelopment of offspring, Nutrition Reviews 76 (2018) 708–721. 10.1093/nutrit/nuy025.

[27] E.M. Moore, D. Ames, A.G. Mander, R.P. Carne, H. Brodaty, M.C. Woodward, K. Boundy, K.A. Ellis, A.I. Bush, N.G. Faux, R.N. Martins, C.L. Masters, C.C. Rowe, C. Szoeke, D.A. Watters, Among vitamin B12 deficient older people, high folate levels are associated with worse cognitive function: Combined data from three cohorts, Journal of Alzheimer’s Disease 39 (2014). 10.3233/JAD-131265.

[28] S. Deghan Manshadi, L. Ishiguro, K.-J. Sohn, A. Medline, R. Renlund, R. Croxford, Y.-I. Kim, Folic Acid supplementation promotes mammary tumor progression in a rat model., PloS One 9 (2014) e84635–e84635. 10.1371/journal.pone.0084635.

[29] C. Bennett, J. Green, M. Ciancio, J. Goral, L. Pitstick, M. Pytynia, A. Meyer, N. Kwatra, N.M. Jadavji, Dietary folic acid deficiency impacts hippocampal morphology and cortical acetylcholine metabolism in adult male and female mice, Nutritional Neuroscience (2021) 1–9. 10.1080/1028415X.2021.1932242.

[30] K.M. Moon, J. Kim, Y. Seong, B.-C. Suh, K. Kang, H.K. Choe, K. Kim, Proprioception, the regulator of motor function, BMB Rep 54 (2021) 393–402. 10.5483/BMBRep.2021.54.8.052.

[31] L. Venkatasubramanian, R.S. Mann, The development and assembly of the Drosophila adult ventral nerve cord, Curr Opin Neurobiol 56 (2019) 135–143. 10.1016/j.conb.2019.01.013.

[32] D. Zhou, G.G. Haddad, Genetic Analysis of Hypoxia Tolerance and Susceptibility in Drosophila and Humans, Annu. Rev. Genom. Hum. Genet. 14 (2013) 25–43. 10.1146/annurev-genom-091212-153439.

[33] G.G. Haddad, R.J. Wyman, A. Mohsenin, Y.-A. Sun, S.N. Krishnan, Behavioral and Electrophysiologic Responses of Drosophila melanogaster to Prolonged Periods of Anoxia, J Insect Physiol 43 (1997) 203–210. 10.1016/s0022-1910(96)00084-4.

[34] P. Azad, D. Zhou, E. Russo, G.G. Haddad, Distinct mechanisms underlying tolerance to intermittent and constant hypoxia in Drosophila melanogaster, PLoS One 4 (2009) e5371. 10.1371/journal.pone.0005371.

[35] W.A. Van Voorhies, Metabolic function in Drosophila melanogaster in response to hypoxia and pure oxygen, Journal of Experimental Biology 212 (2009) 3132–3141. 10.1242/jeb.031179.

[36] F. Sadre-Marandi, T. Dahdoul, M.C. Reed, H.F. Nijhout, Sex differences in hepatic one-carbon metabolism, BMC Systems Biology 12 (2018). 10.1186/s12918-018-0621-7.

[37] P. García-Rodríguez, F. Ma, C.D. Río, M. Romero-Bernal, A.M. Najar, M. de la L. Cádiz-Gurrea, F.J. Leyva-Jimenez, L. Ramiro, P. Menéndez-Valladares, S. Pérez-Sánchez, A. Segura-Carretero, J. Montaner, Diet Supplementation with Polyphenol-Rich Salicornia ramosissima Extracts Protects against Tissue Damage in Experimental Models of Cerebral Ischemia, Nutrients 14 (2022) 5077. 10.3390/nu14235077.

[38] P. Habib, J. Jung, G.M. Wilms, A. Kokott-Vuong, S. Habib, J.B. Schulz, A. Voigt, Posthypoxic behavioral impairment and mortality of Drosophila melanogaster are associated with high temperatures, enhanced predeath activity and oxidative stress, Exp Mol Med 53 (2021) 264–280. 10.1038/s12276-021-00565-3.

[39] I. Celardo, S. Lehmann, A.C. Costa, S.H. Loh, L. Miguel Martins, dATF4 regulation of mitochondrial folate-mediated one-carbon metabolism is neuroprotective, Cell Death Differ 24 (2017) 638–648. 10.1038/cdd.2016.158.

[40] S.L. Helfand, B. Rogina, Genetics of Aging in the Fruit Fly, Drosophila melanogaster, Annual Review of Genetics 37 (2003) 329–348. 10.1146/annurev.genet.37.040103.095211.

[41] D. Amoah, M. Schmidt, C. Mather, S. Prior, M.P. Herath, M.-L. Bird, An international perspective on young stroke incidence and risk factors: a scoping review, BMC Public Health 24 (2024) 1627. 10.1186/s12889-024-19134-0.

